# Contamination of Currency Notes with Kanamycin Resistant *Shigella flexneri*

**DOI:** 10.1101/2020.03.07.982017

**Authors:** Ebrahim Mohammed Al-Hajj, Malik Suliman Mohamed, Noha A. Abd Alfadil, Hisham N. Altayb, Abeer Babiker Idris, Salah-Eldin El-Zaki, Mohamed A. Hassan

## Abstract

*Shigella flexneri* is the main causative agent of shigellosis commonly distributed in developing countries with high morbidity and mortality rates. This study aimed to examine the presence of *Shigella* species in Sudanese currency notes using both traditional and molecular techniques. One hundred thirty five currency notes were collected and their contaminants were isolated and identified conventionally and genetically using 16S rRNA gene amplification and sequencing. Eight isolates were identified as *Shigella* species in different notes, and 3 of them were resistant to penicillin, kanamycin and nitrofurantoin. One *S. flexneri* isolate has insertion mutation of guanine nucleotide at position 730 of life’s essential gene 16S rRNA which known evolutionarily to be stable gene. Banknotes are highly circulating items and therefore, appropriate measures such as regular replacement of the dirty notes with new papers are necessary to protect peoples from being infected with drug resistant pathogens.

## 1. Introduction

*Shigella flexneri* is a gram-negative, non-sporulated, non-motile and anaerobic facultative bacillus bacterium belongs to Enterobacteriaceae family (Penatti *et al*., 2007; Sun *et al*., 2013). *Shigella* species are common pathogenic bacteria which cause shigellosis that is characterized by acute bloody diarrhea (Sansonetti, 2001). The transmission of *S. flexneri* normally occur through fecal-oral route (Jennison and Verma, 2004) and its infections are limited to human only (Sansonetti, 2001; Gaurav *et al*., 2013). Annually, *Shigella spp*. causes 164.7 million infections over the world. In addition, children less than five years old are highly suffering from shigellosis and estimated to be 69% of all *Shigella*s’ infections, while 61% of all deaths due to shigellosis occur in children (Kotloff *et al*.,1999). *Shigella flexneri* is the most frequent cause of shigellosis in developing countries with high morbidity and mortality rates (Kotloff *et al*.,1999; Vinh *et al*., 2009), and it was responsible for many outbreaks in the areas with crowding, poor sanitation and poor hygiene (Walden *et al*., 2005; Ranjbar *et al*., 2010; Shen *et al*., 2017). In 2004, an outbreak of shigellosis (1340 cases and 11 deaths) has been detected in North Darfur (Province located in the western of Sudan) (WHO, 2004). There is a variation in dominance of *S. dysenteriae* and *S. flexneri* in developing countries. Some studies conducted in Khartoum have reported that *S. flexneri* was the most common cause of shigellosis in Sudan (Saeed *et al*., 2015; Ahmed *et al*., 2000). Although shigellosis might be self-limited, but sever cases might be treated with various antibiotics such as ampicillin, co-trimoxazole and nalidixic acid. However, *Shigella flexneri* developed resistance to these drugs, therefore, WHO recommended ciprofloxacin as the drug of choice for treatment of shigellosis and ceftriaxone as an alternative treatment in the last decade (WHO, 2005), which also became ineffective in some cases. *Shigella flexneri* was identified by many methods including conventional microbial methods (Gaurav *et al*., 2013), identification of virulence plasmid with direct PCR assay (Mokhtari *et al*., 2012; Sansonetti *et al*., 1982) and 16S rRNA gene sequencing (Šimenc *et al*., 2008). The 16S rRNA gene, presents in almost all bacteria, is an essential gene for organisms survival (Yıldırım *et al*., 2011), and used in many studies for identification and classification of bacteria at strain level. The gene sequence, approximately 1500 bp, is suitable to contain phylogenetic information and its function is stable for long period of time (Patel, 2001; Janda and Abbott, 2007; Suardana, 2014). In fact, 16S rRNA gene was extensively used in taxonomy and identification of bacteria to species and strain levels (Gee *et al*., 2004; Woo *et al*., 2008), and also used to understand the resistance mechanisms of some bacteria to antibiotics (Springer *et al*., 2001). It is known that currency notes are widely used almost by all nations every day that subjected them to be contaminated by pathogenic microorganisms and transmission of these organisms between people (Gedik *et al*., 2013; Alemu, 2014; Osman *et al*., 2017). *Shigella* species might be one of these organisms that contaminate the currency notes. Therefore, this study was conducted to explore whether *Shigella flexneri* could be detected in Sudanese currency notes with the aid of 16S rRNA gene sequencing, which is more accurate and somewhat consuming less resources and time than conventional methods. In addition, *in silico* tools were used to understand the evolution of *S. flexneri*’s 16S rRNA gene circulating in Sudanese currency notes.

## 2. Materials and Methods

### 2.1 Sample Collection

A total of 135 Sudanese currency notes that randomly collected from hospitals, restaurants, public transportations and banks in Khartoum state were transferred into sterile plastic petri dishes to the laboratory of Pharmaceutical Microbiology in Omdurman Islamic University.

### 2.2 Phenotypic Identification

#### 2.2.1 Isolation and Identification of *Shigella* species

Various microbiological techniques described previously (Cheesbrough, 2006) for bacterial identification were used to identify *Shigella* species. Both sides of currency notes that wetted with sterile distilled water were swabbed using cotton-tipped swab, inoculated on 5% blood agar plates and incubated aerobically at 37 °C for 24 hours. Separated colonies were sub-cultured in MacConkey agar, nutrient agar and xylose lysine desoxycholate agar plates. Biochemical tests: hydrogen sulfide, indole, motility, urease, voges proskauer, methyl red, citrate utilization, glucose fermenter, lactose fermenter, oxidase, catalase tests and gram staining techniques were carried out for preliminary identification of *Shigella* species.

#### 2.2.2 Antibiotic sensitivity Test

Susceptibilities were investigated by disk diffusion method according to the protocol specified previously (Forbes *et al*., 2007), while inhibition zone diameters interpretations were carried out according to the European Committee on Antimicrobial Susceptibility Testing (EUCAST, 2016). Antibiotics used in this study were: amoxicillin (25μg), ampicillin (30μg), co-amoxyclav (30μg), cephalexin (30μg), cefuroxime (30μg), ceftriaxone (30μg), ceftazidime (30μg), gentamicin (10μg), kanamycin (30μg), co-trimoxazole (25μg), penicillin-G (10 IU), erythromycin (5μg), azithromycin (15μg), ciprofloxacin (30μg), levofloxacin (5μg), nitrofurantoin (200μg), vancomycin (30μg), chloramphenicol (30μg) and meropenem (10μg).

### 2.3 Molecular analysis

16S rRNA gene analysis confirmatory test was carried out for the isolates identified as *Shigella* species and showed multiple drug resistance.

#### 2.3.1 DNA Extraction

The bacterial genomic DNA was extracted using Chelex 100 method (Giraffa *et al*., 2000). Briefly, three colonies were transferred to a tube containing 200 μL of 1X PBS, mixed by vortex for ten seconds using a vortex shaker (GEMMY industrial CORP, Taiwan), and centrifuged for five minutes. The supernatant was discarded and the pellet was suspended in 200 μL of 6% Chelex reagent and incubated at 56 °C for fifteen to thirty minutes, and further boiled at 100 °C for fifteen minutes. After ten seconds vortexing and cooling at room temperature, the supernatant was collected with the aid of centrifugation.

#### 2.3.2 16S rRNA gene amplification

The 16S rRNA gene was amplified from genomic DNA by PCR using thermocycler (BIO-RAD, Mexico). Universal primers of 16S rRNA gene 27F (5’-AGAGTTTGATCCTGGCTCAG-3’) and 1495R (5’-CTACGGCTACCTTGTTACGA-3’) were used (Liu *et al*., 2009). The PCR was carried out using PreMix Mastermix kit (iNtRON Biotechnology) in 25 μL reaction mixture containing 13 μL distill water, 1 μL forward primer, 1 μL reverse primer and 5 μL DNA extract. The PCR consists of initial denaturation step at 94 °C for 5 min followed by 30 cycles of [94 °C for1 min, 58 °C for 1 min and 72 °C for 2 min], and final extension step at 72 °C for 10 min (Mokhtari *et al*., 2012). The PCR product was analyzed using 1% agarose gel electrophoresis (Lee *et al*., 2012) to check the presence of amplified fragment, 1500 bp, and sent to Macrogen Company in Netherlands for further purification and sequencing. The obtained sequence was deposited in the GenBank database under accession number KY199565 and the strain was named “ebra1”.

### 2.4 Bioinformatics Analysis

The obtained sequence was manually cleaned by Finch TV software version 1.4.0 (Geospiza, Inc.; Seattle, WA, USA), and used to search the NCBI BLASTN program (Morgulis *et al*., 2008) for similar sequences. The obtained sequence was aligned with the top 15 similar sequences, that found to be of *Shigella flexneri* strains from different countries (Table 1) (Benson *et al*., 2005), using Unipro UGENE software version 1.25.0 (Okonechnikov *et al*., 2012). To study the evolutionary relationships between sequences, phylogenetic tree was constructed by MEGA software version 7 (Kumar *et al*., 2016) using Maximum Likelihood method of analysis (Tamura and Nei, 1993).

**Table 1:**
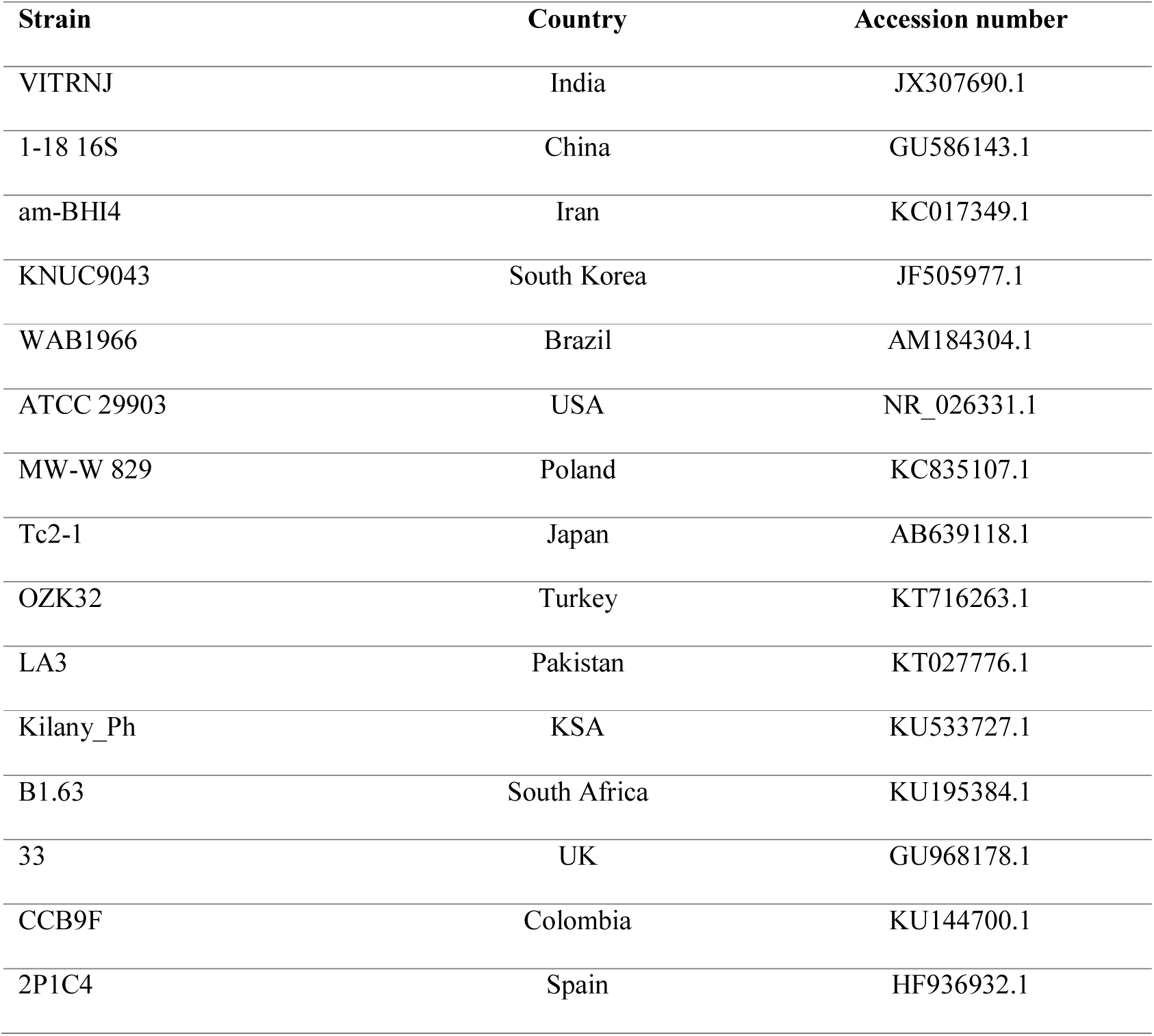
*Shigella flexneri* strains of other countries used in this study.

| Strain | Country | Accession number |
| --- | --- | --- |
| VITRNJ | India | JX307690.1 |
| 1-18 16S | China | GU586143.1 |
| am-BHI4 | Iran | KC017349.1 |
| KNUC9043 | South Korea | JF505977.1 |
| WAB1966 | Brazil | AM184304.1 |
| ATCC 29903 | USA | NR_026331.1 |
| MW-W 829 | Poland | KC835107.1 |
| Tc2-1 | Japan | AB639118.1 |
| OZK32 | Turkey | KT716263.1 |
| LA3 | Pakistan | KT027776.1 |
| Kilany_Ph | KSA | KU533727.1 |
| B1.63 | South Africa | KU195384.1 |
| 33 | UK | GU968178.1 |
| CCB9F | Colombia | KU144700.1 |
| 2P1C4 | Spain | HF936932.1 |

## 3. Results

### 3.1 Isolation and Identification

Eight selected gram negative isolates from different banknotes (5.93% from 135 notes) were identified as *Shigella* species using biochemical tests (Table 2).

**Table 2:** Biochemical test results for 8 isolated colonies.

| Test | H<sub>2</sub>S<sup>a</sup> | Indole | Motility | Urease<br>test | Citrate<br>utilization | VP<sup>b</sup> | MR<sup>c</sup> | Lactose<br>fermenter | Glucose<br>fermenter | Catalase<br>test | Oxidase<br>test |
| --- | --- | --- | --- | --- | --- | --- | --- | --- | --- | --- | --- |
| Result | - | - | - | - | - | - | + | - | - | + | - |

<sup>a</sup> (H<sub>2</sub>S): Hydrogen Sulfide; <sup>b</sup> (VP): Voges Proskauer; <sup>c</sup> (MR): Methyl Red.

### 3.2 Sensitivity Test

The first *Shigella* isolate was found to be resistant to penicillins (amoxicillin, ampicillin, co-amoxyclav and penicillin-G), kanamycin and nitrofurantoin, while the second isolate was found to be resistant to tested penicillins, but not co-amoxyclav. The remaining 6 isolates were found to be sensitive to all antibiotics tested (Table 3).

**Table 3:**
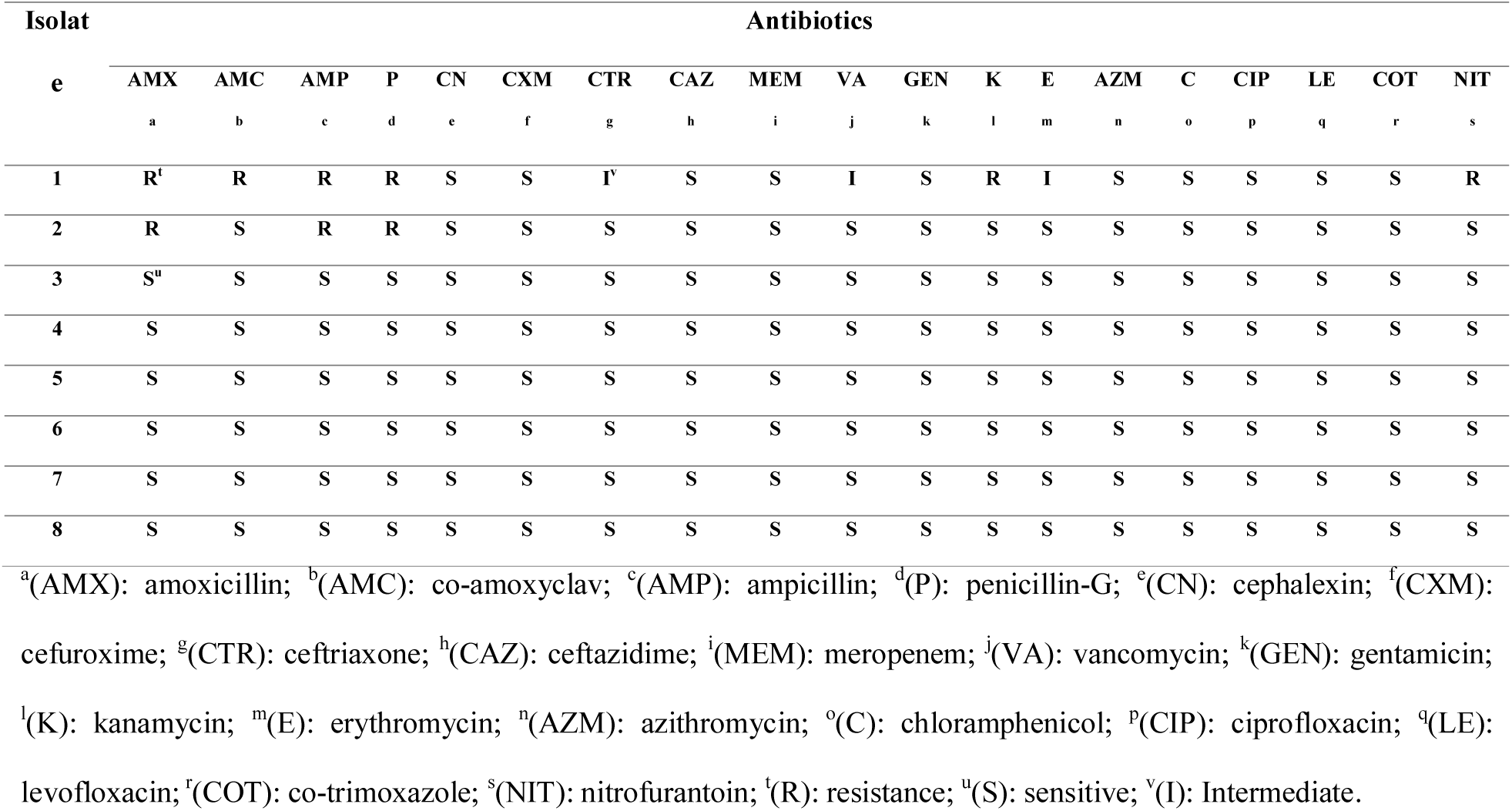
*Shigella* isolates sensitivities.

### 3.3 Bioinformatics Analysis

The 16S rRNA gene sequence of our sample *ebra1*, after cleaning (702 base pairs), was found to be 100% identical to *Shigella flexneri* (strain VITRNJ, India: JX307690.1) based on BLASTN search output. Their alignment to other related sequences including the reference (strain ATCC 29903, USA: NR_026331.1) showed that there is a guanine nucleotide insertion at position 730 (Figure 1.C), and also, they are clustered together in the same phylogenetic tree sub-branch, (Figure 1.D).

**Figure 1:**
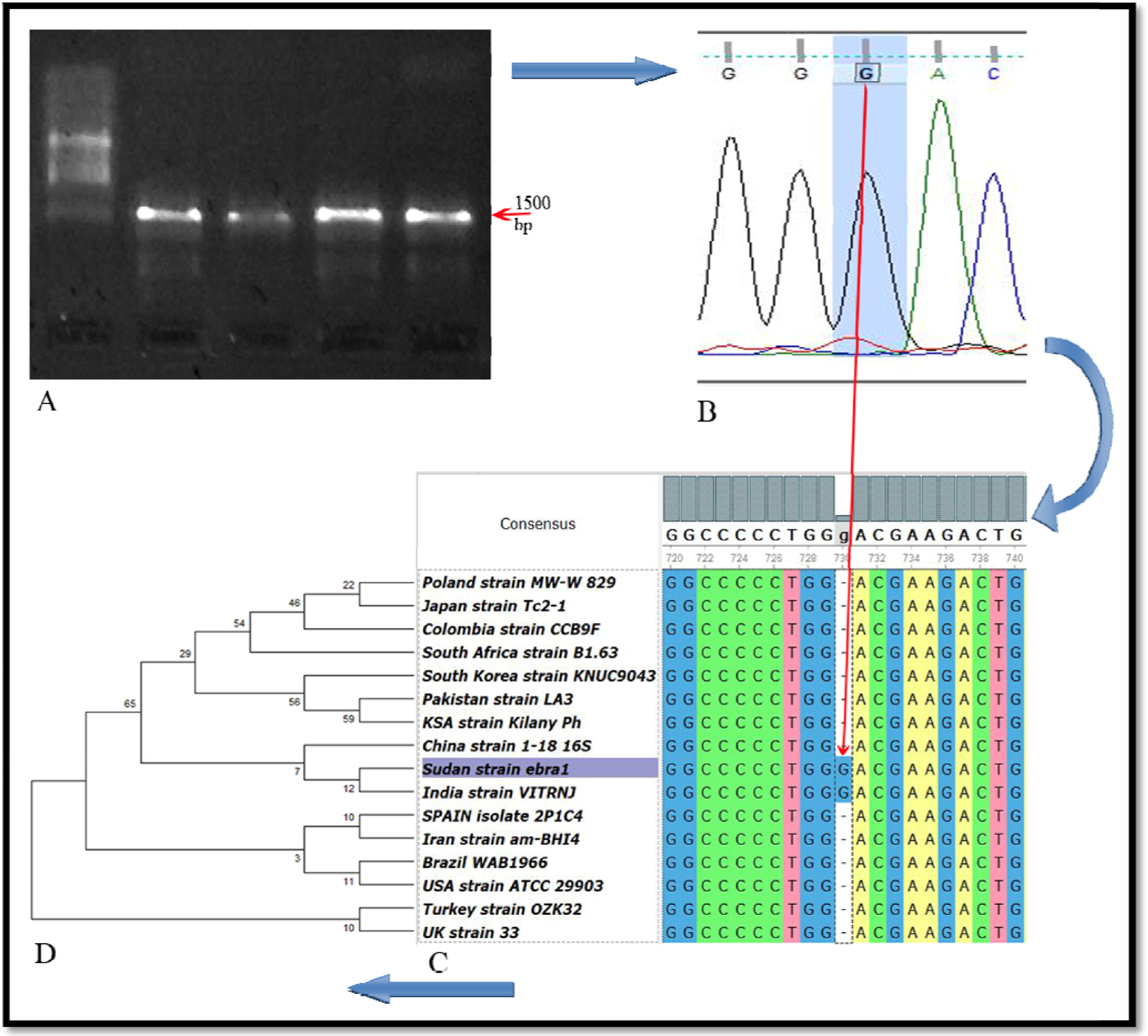
**(A):** PCR product of the *S. flexneri*’s 16S rRNA gene run over % agarose gel electrophoresis. **(B):** 16S rRNA gene sequence chromatogram, shown by FinchTV software, indicating a guanine nucleotide insertion at position 730. **(C):** Alignment of *ebra1* and *JX307690*.*1* to other related sequences determining a guanine nucleotide insertion mutation using Unipro UGENE software version 1.25.0. **(D):** Phylogenetic analysis of 16S rRNA gene sequences of Sudan strain, *ebra1*, and selected strains of *S. flexneri* of other countries using MEGA software version 7.

## 4. Discussion

16S rRNA gene sequencing has become one of the most preferred technique for identification of bacteria that it might give information to species and strain level. We identified *Shigella spp*. in banknotes by phenotypic method and confirmed it as *S. flexneri* using 16S rRNA gene sequencing. There are some difficulties facing researchers identifying bacteria by phenotypic approaches such as the length of the tests and variations of strains in the same species. Many authors prefer partial 16S rRNA gene sequencing method (Šimenc *et al*., 2008) for bacterial identification, which gives more accurate, reproducible and robust results (Clarridge, 2004). *Shigella* is one of the organisms that should not contaminate our daily contact objects such as banknotes, due to the fact that it causes more severe disease with greater morbidity (Navia *et al*., 2005). It is a highly contagious organism with infective dose of only 10–100 viable cells and an incubation period of 1–5 days. Better community hygiene practices could help in reducing the chance of *Shigella* diseases, which transmitted mainly to fomites with contaminated hands and other interpersonal contacts (Ko *et al*., 2013; Li *et al*., 2016). Although *Shigella* is a fragile and remains viable for a limited time outside human body (Sack *et al*., 2001), but we detected *Shigella spp*. in 5.93% of the tested Sudanese currency notes, that might be a potential source of infection. It seems that countries borders do not prevent the transmission of social cultures between nations, this could be reflected partially by this finding and the presence of *Shigella* in Pakistani currency notes (6% from 167) (Ali *et al*., 2015), possibly, the organism from feces transferred to hands and then to currency. There is a possibility that the rate of *Shigella* contamination is even higher than what we found, and using culture-independent approaches for identification may increase the rate. But the identification of drug resistance patterns would not be possible using these approaches (Jalali *et al*., 2015).

Multiple sequence alignment of *ebra1* with the other similar sequences retrieved from GenBank database shows insertion mutation of guanine nucleotide in *ebra1* and JX307690.1 genes in position 730. Actually these 2 genes bearing 100% similar sequences (Figure 1.C), and they are clustered together in the same phylogenetic tree sub-branch (Figure 1.D). While GU586143.1 gene of China strain is exactly similar to these two genes without this insertion mutation; consequently, they were clustered together in the same phylogenetic tree branch. It is not clear that whether this mutation is implicated in drug resistance or not, however, the *S. flexneri* isolate showed resistance to different antibiotics including penicillins (amoxicillin, ampicillin, co-amoxyclav and penicillin-G), kanamycin and nitrofurantoin. Likewise, *S. flexneri* isolated from different samples worldwide such as Sudan (Ahmed *et al*., 2000), India (Hosseini *et al*., 2010; Bhattacharya *et al*., 2014), China, (Pan *et al*., 2006; Pu *et al*., 2009), Taiwan (Ko *et al*., 2013), Nepal (Parajuli *et al*., 2017), Malaysia (Koh *et al*., 2012), Missouri, USA (Arvelo *et al*., 2009), New Zealand (Upton *et al*., 2007), Australia (Stafford *et al*., 2007) and Yemen (Al-Moyed *et al*., 2006) were resistant to several antibiotics, even those commonly used for their treatment. The exact reasons behind spread of resistance might not be fully clear, however, socioeconomic and behavioral factors (Okeke *et al*., 1999), and travelling (Navia *et al*., 2005) might result in increasing the spread of multi-drug resistant microorganisms. The molecular bases for drug resistance in *Shigella spp*. has been extensively studied and part of these studies with the impact of chromosomal mutations shown in (Table 4).

**Table 4:** *S. flexneri* chromosomal mutations and their resulting phenotypes.

| Gene | Mutation | Affect | Reference |
| --- | --- | --- | --- |
| *DNA Gyrase*<br>*(gyrA) gene* | G→T substitution, leading to a | fluoroquinolone resistance | (Chu *et al.*, 1998a; Oonaka |
|  | Asp→Tyr substitution at amino acid |  | K *et al.*, 1998) |
|  | residue 87 |  |  |
|  | Asp→ Asn substitution at amino acid |  | (Pazhani *et al.*, 2008) |
|  | residue 87 |  |  |
|  | C→T | Nalidixic acid resistance | (Li *et al.*, 2016; Oonaka K |
|  | substitution, leading to a Ser→ Leu |  | *et al.*, 1998; Pazhani *et al.*, |
|  | substitution at amino acid residue 83 |  | 2008; Chu *et al.*, 1998b) |
| *Topoisomerase IV*<br>*(parC) gene* | A→C substitution, leading to a Asp→ | fluoroquinolone resistance | (Chu *et al.*, 1998a) |
|  | Ala substitution at amino acid residue |  |  |
|  | 79 |  |  |
|  | A→C substitution, leading to a Glu→ |  | (Chu *et al.*, 1998a) |
|  | Ala substitution at amino acid residue |  |  |
|  | 84 |  |  |
| *bla<sup>OXA-1</sup> gene* | A→G | resulting in the novel | (Siu *et al.*, 2000) |
|  | substitution, leading to a Arg→ Gly | designation OXA-30 in |  |
|  | substitution at amino acid residue 131 | ampicillin-resistant |  |
|  |  | *S.flexner* |  |
| *Glucosyltransferas*<br>*e gtrII gene* | G→A substitution at 3476 bp, leading | Conversion of *Shigella* | (Chen *et al.*, 2003) |
|  | to a Cys→Tyr substitution at amino | *flexneri* serotype 2a to |  |
|  | acid residue 437 | serotype Y |  |

On the other hand, mutations in 16S rRNA gene is not without value, they might result in resistance to protein synthesis inhibitors especially those bind to the 30S subunit of microbial ribosomes as illustrated by Moazed and Noller (Moazed and Noller, 1987), and several other studies shown in (Table 5).

**Table 5:** Mutations in 16S rRNA gene that documented by several species and strains.

| **Mutation** | **Affect** | **Reference** |
| --- | --- | --- |
| **G360A** | tetracycline resistance | (Trieber and Taylor 2002) |
| **triple mutation, AGA<sub>965-967</sub>TTC** |  | (Trieber and Taylor 2002; Gerrit *et al.*, 2003) |
| **deletion of G942** |  | (Trieber and Taylor 2002) |
| **deletion of G771** |  | (Trieber and Taylor 2002) |
| **G1058C mutation** | Drug resistance in *Propionibacterium acnes* | (Ross *et al.*, 1998) |
| **Insertion of C530** | Streptomycin resistance and dependence | (Honore *et al.*, 1995) |
| **C512, A513, and C516 are substituted by other bases** | Streptomycin resistance | (Honore *et al.*, 1995) |
| **A1400G, C1401T, G1483T** | kanamycin resistance | (Suzuki *et al.*, 1998) |
| **A1408G** | amikacin, kanamycin and gentamicin resistance | (Nessar *et al.*, 2001; Prammananan *et al.*, 1998; Recht *et al.*, 1999) |
| **T1406A, C1409T, G1491T, G1449C** | kanamycin resistance and reduction of other aminoglycosides binding affinity | (Nessar *et al.*, 2001; Prammananan *et al.*, 1998; Recht *et al.*, 1999) |

It is known that there are various mechanisms behind aminoglycosides resistance, however, resistance of *S. flexneri* to kanamycin detected in this study might be attributed to the insertion mutation of guanine in 16S rRNA gene that might need further studies to be confirmed. There is no doubt that drug resistance might exaggerate the threat of shigellosis especially in developing countries where shigellosis is endemic (Sack *et al*., 2001).

The presence of multi-drug resistant microorganisms in highly circulating items such as currency notes will be of significant health implications. Improving personal hygiene and raising the public awareness might tackle the spreading of these microbes.

## 5. Conclusions

We have detected a multi-drug resistant strain of *S. flexneri* in currency notes circulating in Sudan using conventional methods of identification and 16S rRNA gene sequencing. The identified insertion mutation, G730 in 16S rRNA gene, does not affect ribosome function and the organism was viable in the tested currency notes. However, it might be a mechanism behind kanamycin resistance that detected in this study, possibly, by shifting the nucleotides of the decoding A site (aminoglycoside binding site) since they are downstream to the insertion mutation site. Regular investigations of the presence of infectious agents in currency notes, replacement of the dirty and/or old currency with new ones combined with improving personal hygiene are necessary to avoid pathogenic microbial transmission through currency notes.

